# Direct observation of branching MT nucleation in living animal cells

**DOI:** 10.1101/613463

**Authors:** Vikash Verma, Thomas J. Maresca

## Abstract

Centrosome-mediated microtubule (MT) nucleation has been well-characterized; however, numerous non-centrosomal MT nucleation mechanisms exist. The branching MT nucleation pathway envisages that the γ-tubulin ring complex (γ-TuRC) is recruited to MTs by the augmin complex to initiate nucleation of new MTs. While the pathway is well-conserved at a molecular and functional level, branching MT nucleation by core constituents has never been directly observed in animal cells. Here, multi-color TIRF microscopy was applied to visualize and quantitatively define the entire process of branching MT nucleation in dividing *Drosophila* cells. The stereotypical branching nucleation event entailed augmin first binding to a “mother” MT, recruitment of γ-TuRC after 16s, followed by nucleation 15s later of a “daughter” MT at a 36° branch angle. Daughters typically remained attached throughout their ~40s lifetime unless the mother depolymerized past the branch point. Assembly of branched MT arrays, which did not require D-TPX2 (Drosophila TPX2) or evident regulation by a RanGTP gradient, enhanced localized RhoA activation during cytokinesis.

## INTRODUCTION

Microtubules (MT) play a central role in many biological processes including cell division, cell movement, and intracellular transport. The efficacy of MT-dependent processes depends on spatio-temporal control of MT nucleation as well as the organization of polymerized MTs into specific 3-dimensional arrays. The centrosome is a major MT nucleation center and MT array organizer. However, the fact that centrosomes are absent from most mammalian oocytes/eggs and many plant species (Budde and Heald, 2003; Clift and Schuh, 2013; Severson et al., 2016) was an early indication of the existence of acentrosomal nucleation pathways. One such acentrosomal pathway is MT-dependent branching MT nucleation, which was initially described in plants where the phenomenon was clearly visualized in cortical MT regrowth assays in a green algae (O. WASTENEYS and E. WILLIAMSON, 1989) and then quantitatively analyzed in *Arabidopsis* cortical interphase MT arrays and Tobacco BY-2 cell-free lysates (Chan et al., 2009; Liu et al., 2014; Murata et al., 2005; Nakamura et al., 2010; Walia et al., 2014).

Centrosome-mediated MT nucleation requires γ-tubulin and its associated proteins known as the γ-tubulin ring complex (γ-TuRc) (Kollman et al., 2010; Moritz et al., 2000). γ-TuRC is not limited to the centrosome as the complex localizes along the length of spindle MTs (Goshima et al., 2008; Zhu et al., 2008) and visualization of plant cortical arrays revealed that daughter MT branches were nucleated by mother-associated γ-TuRC (Murata et al., 2005). The mechanism and molecules responsible for recruiting γ-TuRC to mother MTs was unknown until a genome-wide RNAi screen in *Drosophila* S2 cells identified 5 dim γ-tubulin (DGT2-6) proteins that were required for γ-tubulin localization to spindle MTs, but not centrosomes (Goshima et al., 2007). Dgt2-6 formed a stable complex called augmin that was required for proper spindle assembly and function (Goshima et al., 2007). Subsequently, augmin was determined to be a conserved octameric protein complex containing DGT2-9 subunits in *Drosophila* (Uehara et al., 2009) and in human the following 8 subunits of augmin were identified: Ccdc5 (HAUS1), Cep27 (HAUS2), hDgt3 (HAUS3), C14orf94 (HAUS4), hDgt5 (HAUS5), hDgt6 (HAUS6), UCHL5IP (HAUS7), and Hice1 (HAUS8) (Lawo et al., 2009; Uehara et al., 2009). Hice1/HAUS8/Dgt4 has been shown to bind directly to MTs (Hsia et al., 2014; Wu et al., 2008); while the Dgt3, Dgt5, and hDgt6/HAUS6/Dgt6 subunits of augmin have been reported to bind to γ-TuRC via its subunit, NEDD-1 (Chen et al., 2017; Haren et al., 2006; Luders et al., 2006; Uehara et al., 2009; Zhu et al., 2008). Augmin is functionally well-conserved as depletion of augmin components in various cell types leads to reduction in spindle MT density, chromosome mis-segregation, midzone MT assembly defects, and increased incidence of cytokinesis failure (Hayward et al., 2014; Uehara and Goshima, 2010; Uehara et al., 2016; Uehara et al., 2009; Zhu et al., 2008).

Since the identification of augmin, the interphase cortical MT array in *Arabidopsis* has provided the best system to visualize the central players (MTs, augmin, γ-TuRC) during branching nucleation (Liu et al., 2014; Wang et al., 2018). The formation of augmin-dependent branched MT arrays was also visualized in *Xenopus* egg extracts (Petry et al., 2013) although the postulated steps of branching MT nucleation: 1) augmin binding, 2) recruitment of γ-TuRC to mother MTs, and 3) nucleation of daughter MTs were not directly observed. In the egg extract model, both RanGTP and its downstream target Targeting Protein for Xklp2 (TPX2), which mediate acentrosomal spindle assembly around DNA (Groen et al., 2009; Maresca et al., 2009) have been implicated in promoting branching MT nucleation (Alfaro-Aco et al., 2017; Petry et al., 2013). Interestingly, these factors are unlikely to contribute to branching MT nucleation in interphase cortical MT arrays in plant cells as both RanGTP and plant TPX2 are nuclear during interphase (Vos et al., 2008). There is also a discrepancy between measured branch angles across systems with plants exhibiting larger augmin-mediated branch angles (~40°) than those measured in *Xenopus* egg extracts (Chan et al., 2009; Liu et al., 2014; Murata et al., 2005; Nakamura et al., 2010; Petry et al., 2013; Walia et al., 2014). Present understanding of the branching MT nucleation pathway in animal cells is limited due to the absence of direct high-resolution imaging of daughter MT nucleation events by its molecular mediators. In this study, multi-color, live-cell TIRF imaging in *Drosophila Melanogaster* S2 cells, the system in which augmin was first identified (Goshima et al., 2008; Goshima et al., 2007), was applied to visualize the entire process of branching MT nucleation by augmin and γ-TuRC.

## RESULTS AND DISCUSSION

In a prior study, we observed the assembly of astral-like MT arrays that appeared to be generated by branching MT nucleation after anaphase onset in *Drosophila* S2 cells (Verma and Maresca, 2019). To test if these arrays were generated by bona-fide branching MT nucleation events, a stable cell line co-expressing EGFP-α-tubulin and γ-tubulin-Tag-RFP-T was created and imaged with dual color, high-resolution TIRF microscopy. Centrosomal and spindle MT populations of γ-tubulin were observed throughout mitosis and the localization of γ-tubulin puncta to spindle MTs, while abundant, was transient and dynamic. Prior to anaphase onset, individual MTs could not be readily visualized by TIRF microscopy; however, post anaphase onset more stable (relative to pre-anaphase) astral MTs entered the TIRF field where they could be imaged for extended durations (often minutes). Indeed, γ-tubulin puncta were observed to dynamically localize to astral MTs and to reside at the sites of daughter MT nucleation (**Figure 1A, B; Movie 1**). Quantification of a number of bona-fide branching nucleation events revealed that the lag-time between the localization of a γ-tubulin puncta to a mother MT and nucleation of a daughter MT was 15.9s ± 8.8s (Mean ± S.D., n = 41 events) (**Figure 1C**). In contrast to branching MT nucleation events in *Xenopus* egg extracts where reported branch angles were shallow (< 10° for 52% branching events) (Petry et al., 2013); the branching nucleation events observed in anaphase *Drosophila S2* cells exhibited a mean branching angle of 35.7° ± 8.8° (n = 87) and in the same orientation relative to the mother MT (**Figure 1D**). In most cases, as long as the mother MT did not depolymerize past the branch point, the daughter remained attached to the mother MT and exhibited dynamic instability. While variable, the mean lifetime of a daughter MT was 36.0s ± 25.3s (n = 90) (**Figure 1E**) and, over a typical lifetime, a dynamic daughter MT polymerized microns from the branch site (**Figure 1B**). Branches could nucleate along the length of mother MTs, and, in some cases, branching MT nucleation of “granddaughter” MTs from daughters was observed; however, split-branching of growing MT plus ends (Basnet et al., 2018) was not evident.

**Figure 1.**
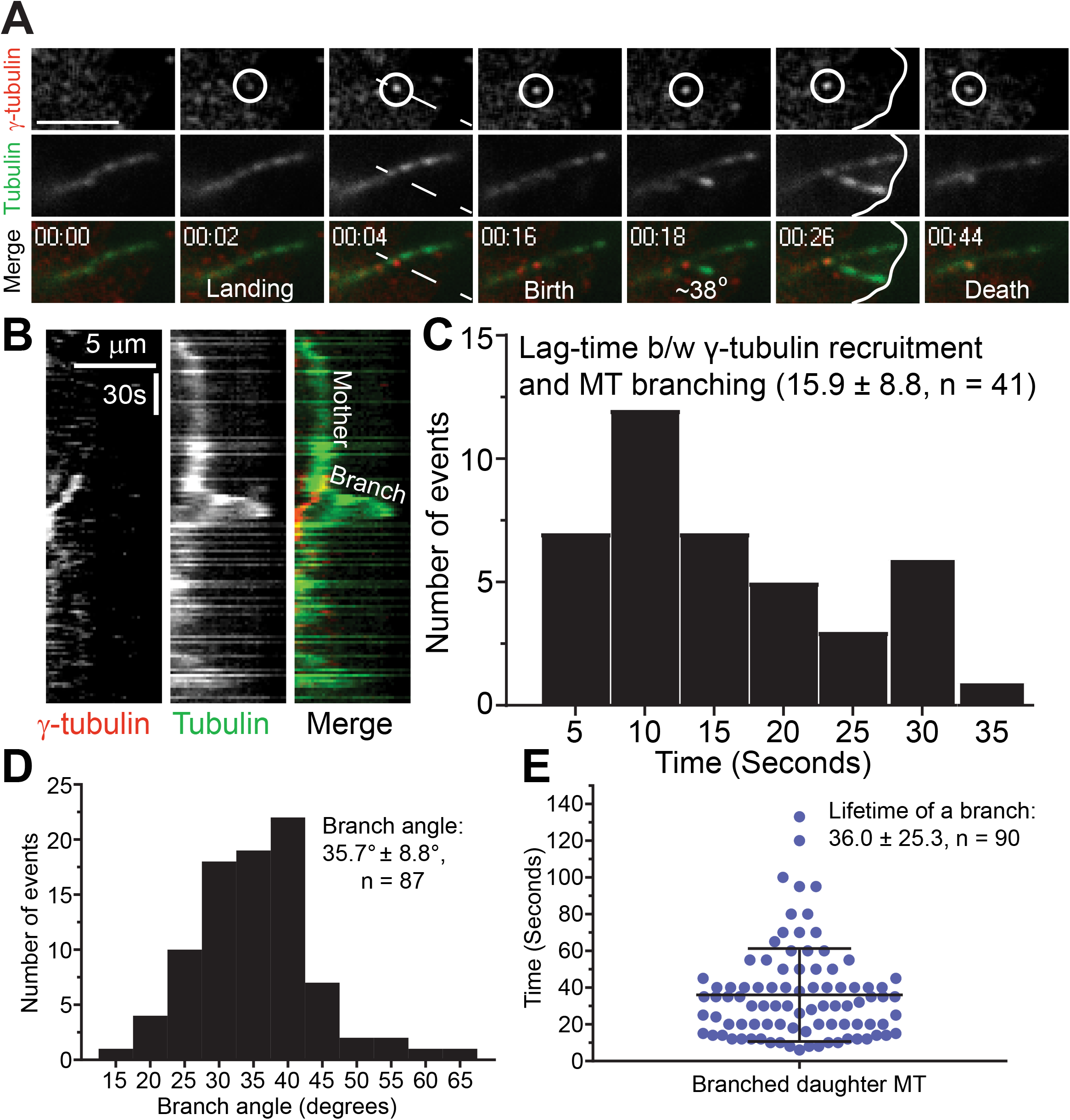
Direct visualization of branching MT nucleation in *Drosophila* S2 cells. (A) Two-color TIRF imaging of an astral MT in a post-anaphase cell co-expressing EGFP-α-tubulin (green), and γ-tubulin-Tag-RFP-T (red). This representative branching event encompasses γ-tubulin landing on the mother MT through disassembly of the daughter MT, both of which contact the cortex (white line at 00:26). (B) Kymograph of the branching event from region (dashed line at 00:04) in Figure 1A. (C) Histogram of the lag-time between γ-tubulin recruitment to the mother MT and birth of a daughter MT, n = 41. (D) Distribution of branch angles between mother and daughter MTs. The branch angles were measured 5-10s after the appearance of daughter MTs, n = 87. (E) Dot plot shows distribution of lifetime of daughter MTs, n = 90, lifetime is time from birth to death of the daughter MT. Time: mins:secs. Scale bars, 5 μm (A, B). Mean ± SD values are reported in all the histograms. Error bars on the dot plot indicate ± SD.

The 36° branch angle measured in S2 cells is entirely consistent with branching MT nucleation in cortical interphase MT arrays in plants (Chan et al., 2009; Liu et al., 2014; Murata et al., 2005; Walia et al., 2014). Multiple studies have reported that ~ 80% of events branched at 40° while ~ 20% branched parallel to the mother MT. While the occurrence of shallow angle/parallel branching events cannot be ruled out, we could not confidently identify bona-fide parallel branch events, as defined by daughters originating from a mother-associated γ-TuRC, because parallel MTs were bundled during cytokinesis. It should be noted that augmin depletion in *Arabidopsis*, resulted in a majority of nucleation events (63%) becoming parallel (Liu et al., 2014) indicative of the 40° MT branch angle being mediated by the augmin complex.

It has been proposed that the augmin complex targets γ-TuRC to pre-existing MTs, which in turn, initiates branching MT nucleation; however, this theoretical series of events has never been directly observed in animal cells. To visualize the properties of the augmin complex and γ-TuRC throughout the branching MT nucleation process in *Drosophila* S2 cells, we made a stable cell line co-expressing EGFP-α-tubulin, γ-tubulin-Tag-RFP-T, and mTurquoise2-Dgt5 (an augmin subunit) and performed multi-color, live-cell TIRF microscopy. Dgt5 localized as discreet puncta to spindle MTs (**Figure 2A**) throughout mitosis. Dgt5 puncta were dynamic but typically remained localized to MTs longer than γ-tubulin puncta and, while bound, Dgt5 often displayed minus end directed movements, albeit many of these events were due to poleward movement of the MTs on which the puncta localized. Three-color TIRF imaging of Dgt5 (augmin complex marker), γ-tubulin (γ-TuRC marker), and α-tubulin revealed the existence of two types of mother MT association events. In ~23% of cases, the Dgt5 and γ-tubulin localized to the mother MT simultaneously – presumably as a pre-assembled complex **Supplementary Figure 1, Movie 2**). In a majority of cases (~77%); however, Dgt5 localization preceded γ-tubulin on the mother MT (**Figure 2B, Movie 3**). Thus, while pre-loaded branching nucleation complexes (augmin + γ-TuRC) likely exist, the augmin complex typically associated with the mother MT first and recruited γ-TuRC from the cytosol to nucleate daughter MTs. When Dgt5 preceded γ-tubulin, the lag-time between augmin complex binding and γ-TuRC recruitment was 14.8s ± 8.9s (n = 21) (**Figure 2C**) while the time between Dgt5 binding to the mother MT, which includes both the pre-loaded (23%) and recruitment (77%) modes, and daughter nucleation was 27.0s ± 20.5s (n = 27) (**Figure 2D**). The Dgt5 and γ-tubulin remained at the branch point throughout the lifetime of the daughter unless the mother MT depolymerized past the branch point. Upon complete depolymerization of the daughter MT, the γ-tubulin puncta at the branch point dissociated from the mother MT (**Figure 1B**) while the augmin complex often remained and was even capable of supporting another round of branching MT nucleation (**Figure 2B; Movie 3**).

TPX2 is a highly conserved protein and its activity has been associated with multiple roles in mitosis, such as, spindle assembly (Gruss et al., 2001; Gruss et al., 2002; Ma et al., 2010; Wadsworth, 2015; Wittmann et al., 2000), MT nucleation, and MT stabilization (Alfaro-Aco et al., 2017; Groen et al., 2009; Petry et al., 2013; Reid et al., 2016; Roostalu and Surrey, 2017). In *Xenopus laevis* egg extracts branching MT nucleation events were observed following the addition of 10-20 fold molar excess TPX2 and RanGTP and branching MT nucleation was not observed in extracts depleted of endogenous TPX2 (Petry et al., 2013). One model of augmin-mediated branching MT nucleation proposes that γ-TuRC is recruited to augmin complex via TPX2 due to the fact that the three components co-immunoprecipitated from egg extracts (Petry et al., 2013). TPX2 contains a composite binding sequence that shows resemblance to the yeast Spc110/Pcp1 motif (SPM) and centrosomin motif (CM1)/γ-TuRC nucleation activator motif (γ-TuNA), deletions of which resulted in reduced or no branching MT nucleation in the *Xenopus* egg extract branching assay (Alfaro-Aco et al., 2017).

**Figure 2.**
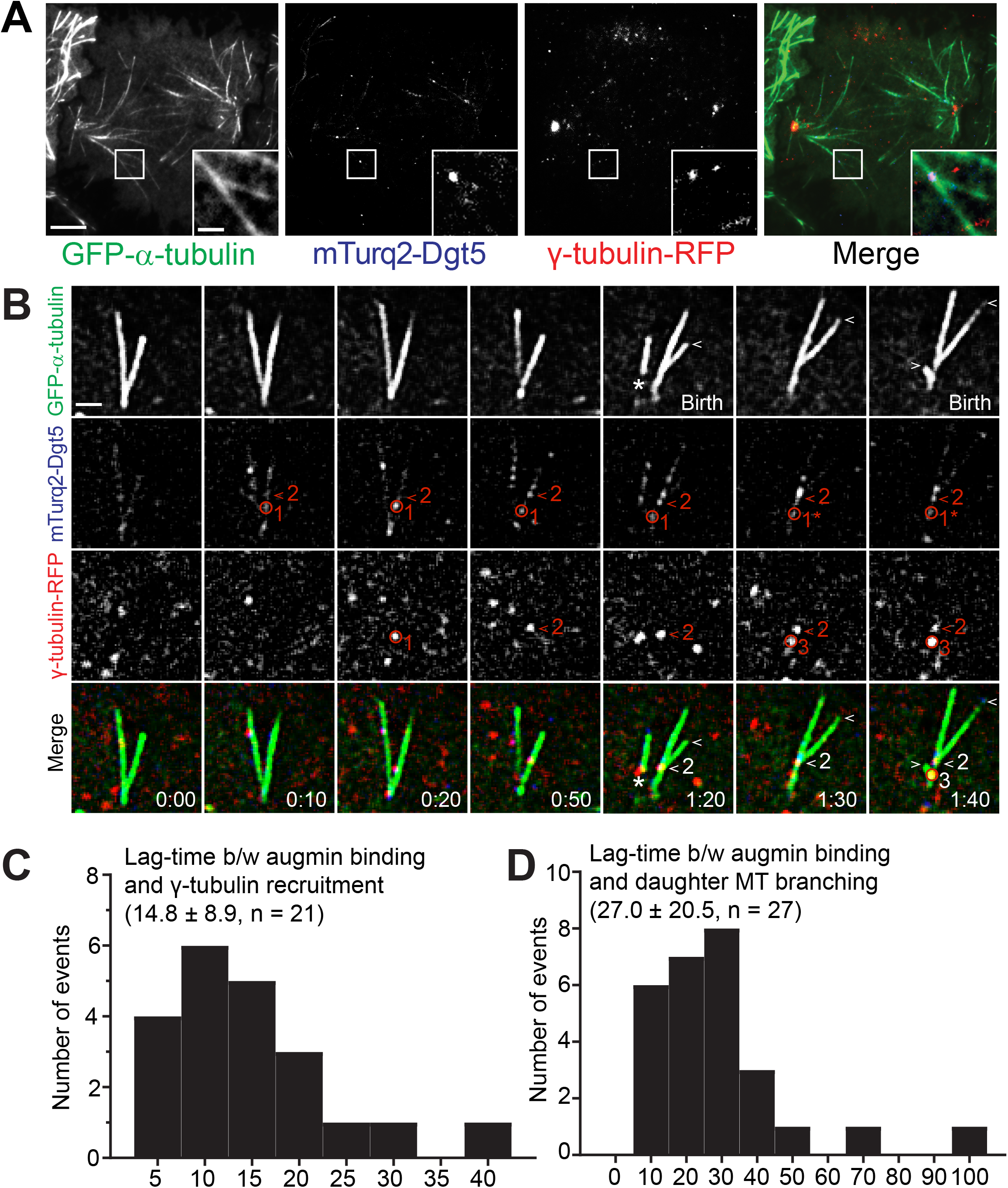
Direct visualization of the key mediators of branching MT nucleation. (A) Representative TIRF micrographs showing expression and localization of γ-tubulin-Tag-RFP-T (red) and mTurquoise2-Dgt5 (blue) in a mid-anaphase *Drosophila* S2 cell. Inset shows a representative MT branching event where mTurquoise2-Dgt5 and γ-tubulin-Tag-RFP-T co-localized. (B) Still-frames from a TIRF time-lapse of MT branching events in a cell co-expressing GFP-α-tubulin (green), mTurquoise2-Dgt5 (blue), and γ-tubulin-TagRFP (red). The events include: 1) localization of two Dgt5 puncta at 10s on mother MT (one indicated by the red circle and marked 1, another indicated by the red arrow head and marked 2 in the mTurquoise2 channel), 2) Dgt5 puncta (1) recruits γ-tubulin (1) (RFP channel, 0:20). The γ-tubulin puncta (1) dissociates from Dgt5 (1) within 30s without a branch event; although, the Dgt5 puncta (1) remains associated with the mother. 3) Dgt5 (2) recruits γ-tubulin (2) (RFP channel, 0:50) and this complex nucleates a daughter within 30 s (1:20). 4) Dgt5 puncta (now denoted 1*) recruits a second γ-tubulin (3) (RFP channel, 1:30), which nucleates a branch within 20s (1:40). The first daughter MT grows for 20 seconds (plus-end indicated by white arrow heads in GFP channel) while the second daughter MT was born shortly before the mother depolymerizes. The asterisk (GFP channel, 1:20) indicates a rare event in which a daughter with minus-end associated γ-tubulin and Dgt5 dissociates from an intact mother. The unattached daughter depolymerized completely within 20s. (C) Average lag-time between mTurquoise2-Dgt5 (augmin complex subunit) binding to the mother MT and recruitment of γ-tubulin-TagRFP-T, n = 21. Binding of pre-assembled complexes of Dgt5 and γ-tubulin, which occurred 23% of the time, are excluded from the histogram. (D) Histogram of lag-time between Dgt5 recruitment to the mother MT and birth of a daughter MT, n = 27. Time: mins:secs. Scale bars, 5 μm (A), 1 μm (A-inset, B). Mean ± SD values are reported.

Sequence alignment between *Xenopus laevis (Xl)* TPX2 and *Drosophila melanogaster (Dm)* D-TPX2 indicated that D-TPX possesses a partially conserved CM1/γ-TuNA motif “ERRRDD’’ while the other conserved motif, SPM, is absent (**Figure 3A**). Further investigation into the predicted secondary structure of D-TPX2 revealed that it’s C-terminus (190-325 amino acids) contains 5 short stretches of α-helices; and the partially conserved γ-TuNA motif lies within the predicted α-helices, as is the case for *Xenopus laevis* (**Figure 3B**) (Alfaro-Aco et al., 2017). To further investigate the role of D-TPX2 in branching MT nucleation events in *Drosophila* S2 cells, we depleted D-TPX2 using RNAi and performed live-cell TIRF microscopy in stable cell lines expressing mTurquoise2-Dgt5, γ-tubulin-Tag-RFP-T, and EGFP-α-tubulin. Consistent with previous reports from live S2 cells, spindle assembly was largely unperturbed following D-TPX2 depletion (> 95%, **Figure 3C**) (Goshima, 2011). Therefore, we were able to readily analyze the effects of D-TPX2 depletion on branching MT nucleation events post-anaphase onset. D-TPX2 depletion did not alter any major effects of TPX2 depletion on MT-dependent branching MT nucleation (**Figure 3D-F**). We measured the same branch angle (**Figure 3E**, Ctrl. RNAi: 36.2° ± 10.4°, n = 22; TPX2 RNAi: 35.7° ± 9.7°, n = 22) and lifetime of daughter MTs (**Figure 3F**. Ctrl. RNAi: 37.7 ± 19.8 Sec, n = 22; TPX2 RNAi: 47.2 ± 20.5 Sec, n = 22) in control and D-TPX2 depleted cells. The branch frequency, as defined by the number of branching events observed in the TIRF field over a 3-minute imaging window post-anaphase onset, was also indistinguishable between control and TPX2-depleted cells. (Ctrl. RNAi: branch/1.3 min; TPX2 RNAi: branch/1.5 min, n = 44 branching events in control and TPX2 RNAi cells). Taken together, these results indicate that D-TPX2 is dispensable for branching MT nucleation in *Drosophila* despite D-TPX2 possessing a putative CM1/γ-TuNA-like motif.

**Figure 3.**
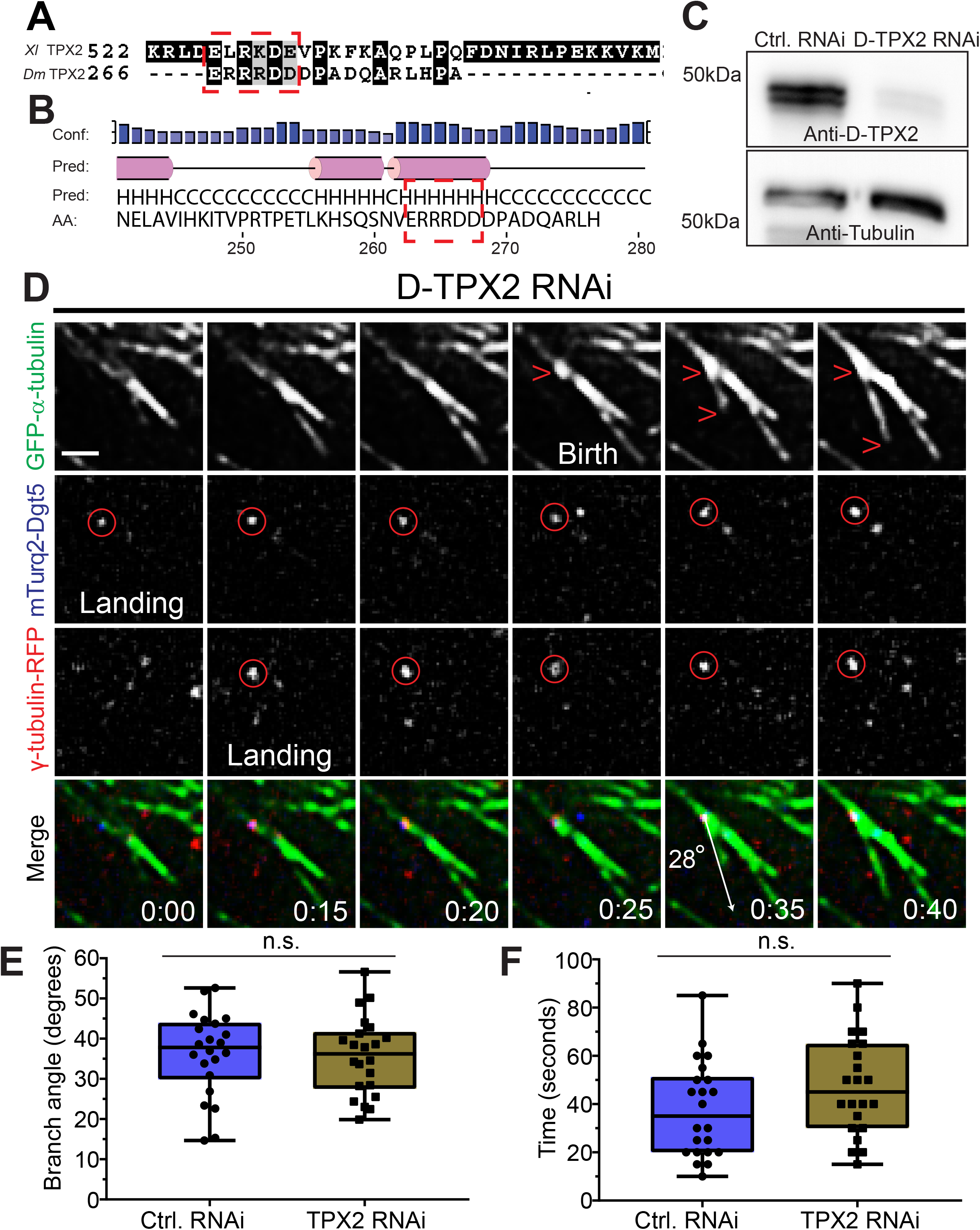
Branching MT nucleation is unaffected by D-TPX2 depletion. (A) Protein sequences of *Xenopus laevis (Xl)* TPX2 and Drosophila melanogaster (*Dm*) TPX2 (D-TPX2) were aligned with T-Coffee multiple alignment software. Red dashed rectangle indicates a partially conserved CM1/γ-TuNA motif in D-TPX2. (B) Secondary structure prediction of D-TPX2 was generated using PSIPRED bioinformatics software. Partially conserved CM1/γ-TuNA motif is expected to lie within α-helices (indicated with red dashed square). (C) Western blot showing depletion of endogenous TPX2 with tubulin as a loading control. (D) A representative MT branching event in a D-TPX2 depleted cell co-expressing GFP-α-tubulin (green), mTurquoise2-Dgt5 (blue), and γ-tubulin-TagRFP (red). Timepoint 00:00 indicates landing of a Dgt5 molecule, which recruits γ-tubulin at 0:15. The Dgt5-γ-tubulin complex nucleates a MT branch at 0:35. (E) Distribution of branch angles in control and D-TPX2 depleted cells, n = 44. (F) Lifetime of branched daughter microtubules in control and D-TPX2 depleted cells, n = 44. Time: mins:secs. Scale bar, 1 μm. Box plots indicate full range of variation (from min to max), the interquartile range and the median. Two-tailed p-values of Student’s t-tests are reported, n.s. is not significant or p > 0.05.

While TPX2 has been proposed to be important for branching in *Xenopus* egg extracts, this observation in *Drosophila* S2 cells is not entirely surprising since: 1) there is a strong effect of augmin depletion on spindle MT density and morphology in S2 cells (Goshima et al., 2008; Goshima et al., 2007), but 2) spindle assembly is largely unperturbed in D-TPX2 depleted cells (Goshima, 2011). If TPX2 directly contributes to branching MT nucleation in vertebrates, then it is conceivable that other factors fulfill the role of TPX2 in *Drosophila*. The RanGTP gradient that is generated around mitotic chromatin (Kalab and Heald, 2008) has also been implicated in promoting branching MT nucleation (Petry et al., 2013). We propose that RanGTP is not required for branching MT nucleation in *Drosophila* S2 cells because: 1) branching nucleation occurs at a significant distance (>5 microns) from segregating chromosomes in anaphase and 2) robust branching nucleation continues throughout telophase after the nuclear envelope has reformed. In light of these observations, it would be worthwhile to investigate the requirement of RanGTP and TPX2 in MT-dependent branching MT nucleation in other model systems.

Augmin-dependent branching MT nucleation contributes to furrow ingression and abscission during cytokinesis (Uehara et al., 2016). To better understand how augmin-dependent branching MT nucleation may contribute to cytokinesis, we performed live-cell microscopy on dividing cells co-expressing Tag-RFP-T-α-Tubulin and EGFP-Rhotekin, a reporter for RhoA-GTP (Bement et al., 2005; Benink and Bement, 2005). During anaphase, we often observed amplification of Rhotekin in the vicinity of large branched MT arrays (**Figure 4A, 4B, Movie 4**). Quantification of Rhotekin fluorescence intensity revealed that localized RhoA-GTP increased by ~ 30% proximal to the mother and daughter MTs during the branching MT event while a nearby cortical area where astral MTs were absent exhibited no change in Rhotekin fluorescence during the same time period (**Figure 4C, 4D**). We propose that localized amplification of RhoA-GTP by branched MT arrays is a consequence of the production of new MT plus-ends that physically contact the plasma membrane, which we recently reported were able to activate cortical RhoA through recruitment of the RhoGEF ECT2 (Verma and Maresca, 2019). In the absence of augmin, compromised midzone assembly has been attributed as the cause of the observed cytokinesis defects (Uehara et al., 2016; Uehara et al., 2009), although we proposed that reduced RhoA activation by branching astral MTs could also contribute to the augmin depletion phenotype.

**Figure 4.**
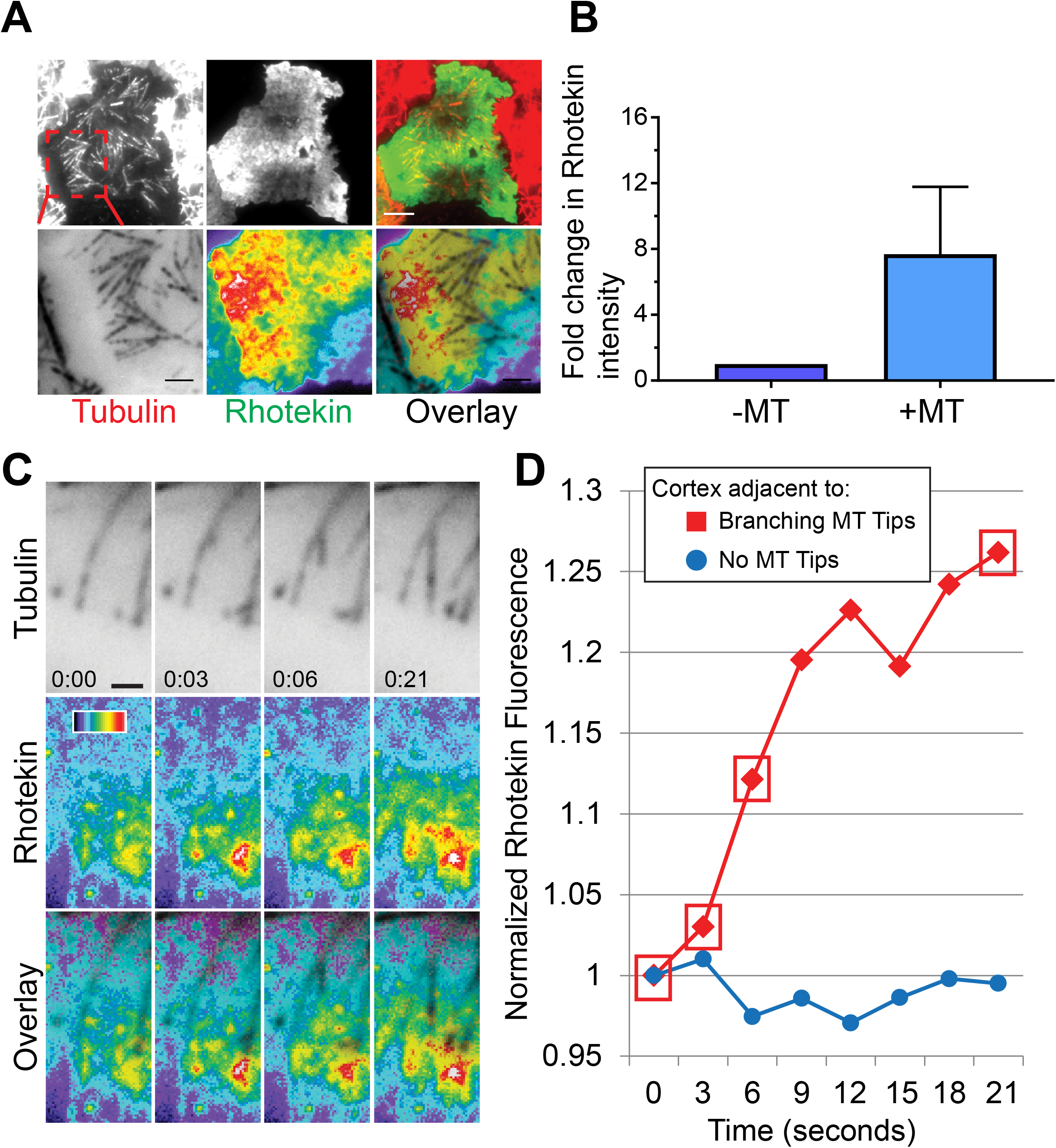
Branching MT nucleation amplifies RhoA activation during cytokinesis. (A) The active RhoA reporter Rhotekin is enriched near the MT plus-ends of branched MT arrays (top); Enlarged views of the region indicated in the inset (red dashed box) of the branched MT array that activates cortical RhoA (bottom). (B) Fold change in Rhotekin fluorescence intensity near the MT plus-ends with respect to a nearby cortical region devoid of astral MTs, data were pooled from 3 independent experiments (n = 3). (C) Still-frames from a multi-color TIRF time-lapse showing a branching event that locally increases cortical RhoA activation - visualized with Rhotekin. (D) Rhotekin fluorescence increases ~30% (red) proximal to the mother and daughter MTs during the branching nucleation event shown in (C). A nearby cortical region devoid of astral MTs (blue) exhibited no change in Rhotekin fluorescence during the same time period. Red boxes correspond to the time points in (C). Error bars indicate SD. Time: mins:secs. Color wedge, pixel values 150-1200. Scale bars, 5 μm (A) 1 μm (C).

The lag-time from γ-tubulin binding to nucleation (15.9. ± 8.8s) and the branch angle (35.7 ± 8.8°) in anaphase *Drosophila* S2 cells are nearly identical to branching parameters quantified in plants (Chan et al., 2009; Liu et al., 2014; Murata et al., 2005; Nakamura et al., 2010; Walia et al., 2014). Interestingly, smaller (< 30°) branch angles have been measured in *Xenopus laevis* egg extracts (Petry et al., 2013), observed by electron tomography of metaphase spindles in human U2OS cells, and inferred from tracking EB3 comets in human HeLa and RPE1 cells (David et al., 2018). This discrepancy may stem from fundamental mechanistic and/or structural differences between the fly/plant and vertebrate branching pathways. However, we favor an alternative explanation for the observed differences in branch angles that are a consequence of two considerations: 1) the structural organization and physical properties of the augmin complex, and 2) the cellular environment in which a daughter is born.

Negative stain EM of the reconstituted octameric complex revealed that human augmin is a ~40-45 nm long, Y-shaped structure with a ~30 nm long stem and ~15 nm flexible splayed end that can adopt multiple conformations (Hsia et al., 2014) (**Figure 5A**). Structural analyses of a human augmin sub-complex indicated that one end of a stem-like structure contains the MT binding region (MTBR) of Hice1/HAUS8 (Hsia et al., 2014). It is compelling to postulate that the splayed Y-shaped end of the complex may contain γ-TuRC binding activities. In sum, the augmin complex may contain a rigid 30 nm long stem that binds MTs at one end and a flexible splayed end that recruits γ-TuRC. Further angular flexibility could be achieved via a flexible hinge - analogous to a wrist - between the MT-binding “arm” and γ-TuRC-binding “hand” (**Figure 5A, B**). Interestingly, negative stain EM of the *Xenopus* augmin holo-complex revealed a stem-like structure bound at a nearly perpendicular angle to the MT with a fair number of MT-bound particles exhibiting hinge-like bends (Song et al., 2018).

**Figure 5.**
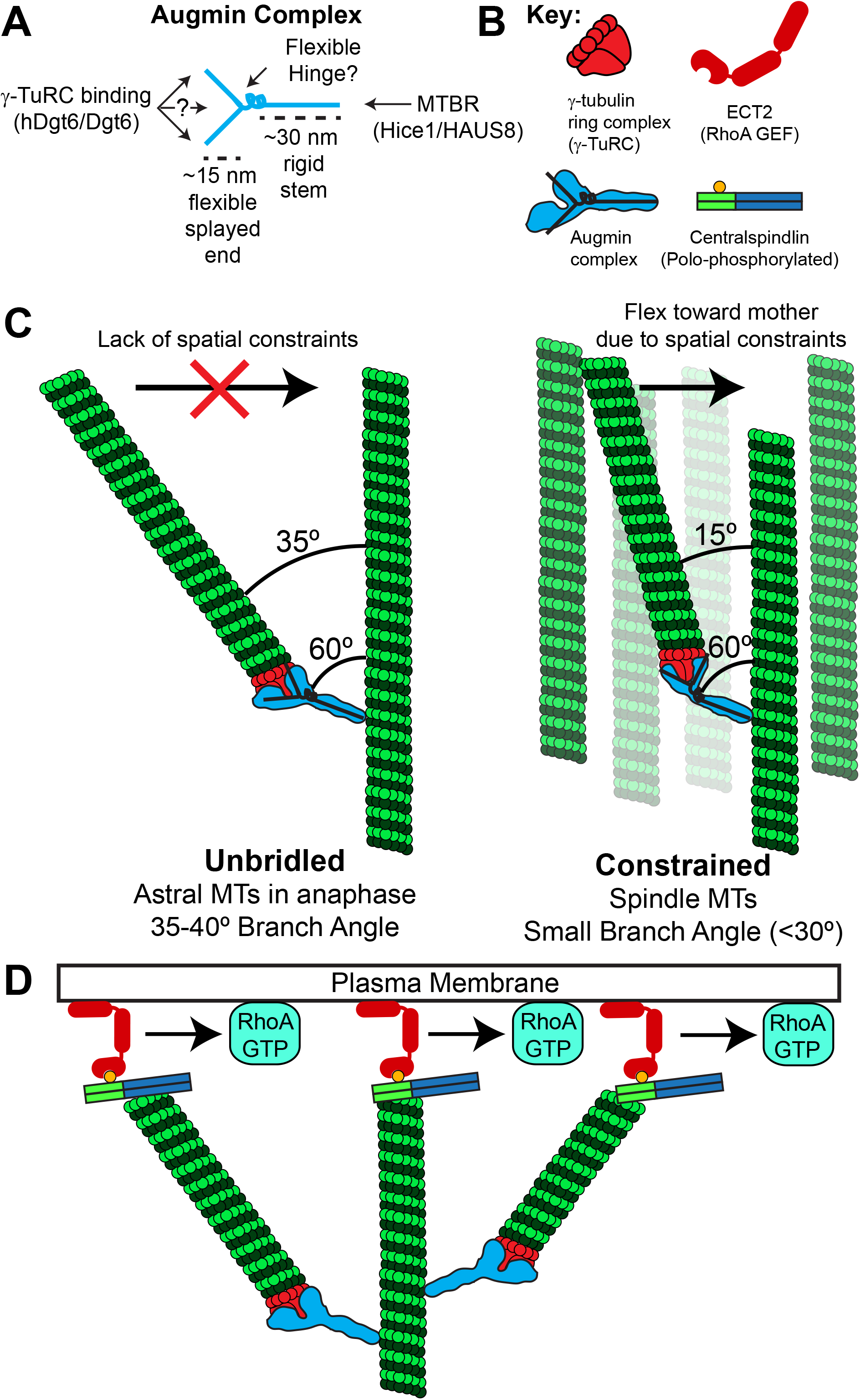
Model of augmin-mediated branching MT nucleation and its role in cytokinesis. Based upon negative stain EM of the augmin complex; we propose that the augmin complex contains a rigid ~ 30 nm long stem that binds MTs at one end; and a ~15 nm flexible splayed end that recruits γ-TuRC through Dgt6 and possibly other interfaces. MTBR = microtubule binding region in Hice 1/HAUS8. (B) Key to molecular schematics in the figure. Note the overlaid “Y” on the augmin complex schematic, which is drawn based on negative stain EM of the complex. (C) A flexible hinge region in the augmin complex and/or flexibility in the splayed Y-end of augmin may allow a daughter MT to sample a broad range of possible branch angles relative to its mother. The branch angle is impacted by the local cellular environment such that “unbridled” daughter MTs with less spatial constraints (e.g. astral MTs in anaphase) would have larger branch angles than daughters nucleated in the spatially “constrained” environment of the spindle and midzone MT array. (D) Branched MT arrays may amplify RhoA activation by generating more MT plus-ends that are capable of recruiting cortical ECT2 via direct interaction with plus-end bound (polo-phosphorylated) centralspindlin.

Augmin complex flexibility would allow a daughter MT to sample a broad range of possible branch angles relative to its mother. We posit that this angle is ultimately dictated by the local cellular environment such that MTs born into “open” cytosol with less spatial constraints, as is the case for the equatorial astral MTs in early anaphase observed here, will have larger branch angles than daughters nucleated in the spatially constrained environment of the spindle and midzone MT array (**Figure 5C**). We envision that “unbridled” daughters nucleated in the absence of major spatial constraints would exhibit a branch angle closer to the angle at which an augmin complex binds to the mother MT. On the other hand, a daughter that encounters spatial constraints as it grows, for example in a spindle with many MTs oriented nearly parallel to the mother, would flex inward due to flexible or hinge-like parts of the augmin complex. Interestingly, mother and daughter MTs of small angle branches visualized by electron tomography in human metaphase spindles were linked by 29 nm “rods” that bound the mother at a 60° angle (Kamasaki et al., 2013). While the molecular identity of these rods was not determined, we agree with the authors’ speculation that the rods may be augmin – specifically the 30nm stem of the augmin complex (Kamasaki et al., 2013).

We propose that branch angle flexibility may allow the branching MT nucleation pathway to make diverse functional contributions throughout mitosis. During spindle assembly and metaphase, shallow branch angles would contribute to the establishment and maintenance of kinetochore-MT attachments. However, the functional contribution of branching is likely different in the context of pioneering astral MTs during anaphase/telophase. We recently characterized a MT-based RhoA activation pathway that functions during anaphase, telophase, and cytokinesis via recruitment of the RhoA GEF ECT2 to cortical sites contacted by MT plus-ends enriched with the centralspindlin complex (Verma and Maresca, 2019) (**Figure 5B, D**). Interestingly, larger branch angle (> 35°) MT arrays produced more plus-ends that contacted the cortex, thereby amplifying localized RhoA activation (**Figure 4A, 5D**). We posit that larger branch angle nucleation from equatorial astral MTs would allow for the activation of cortical RhoA over microns - the size-scale in which a cleavage furrow is initiated and an area significantly larger than a sub-micron sized kinetochore. Furthermore, auto-catalytic amplification of astral MT arrays (Ishihara et al., 2016) via larger branch angle nucleation would more efficiently (relative to shallow branching) fill the MT array expansion volume observed in large cells (100 – 1000 μm) during early embryonic cell divisions (Ishihara et al., 2014; Mitchison et al., 2012; Wuhr et al., 2008). Unlike actin network organization that uses distinct nucleation mechanisms and molecules for branched versus parallel arrays (Svitkina, 2013), angular flexibility of branching MT nucleation may allow a single complex to effectively assemble both branched and near parallel MT arrays.

Over thirty years ago, Salmon and colleagues demonstrated the power of directly observing a cellular phenomenon and quantifying its parameters by visualizing MT dynamic instability *in vitro* and in cells (Cassimeris et al., 1988; Walker et al., 1988). This work transformed thinking around MT dynamics and laid the foundation for decades of research that continues today (Heald and Khodjakov, 2015). To date, the branching MT nucleation pathway has been visualized and quantified in plants but not in animals – a significant knowledge gap given the central importance of this pathway to animal cell division (Lawo et al., 2009; Uehara et al., 2016; Uehara et al., 2009). Much like the quantification of MT dynamic instability parameters accomplished many years ago, we hope that the direct observation of branching MT nucleation and quantification of its key parameters in living animal cells reported here will inform future mechanistic models of cellular processes that depend on dynamic MTs.

## MATERIALS AND METHODS

### DNA constructs

The γ-tubulin gene (CG3157) with its endogenous promoter was amplified from the genomic DNA of *Drosophila*. The resulting PCR product was cloned between 5’ KpnI and 3’ EcoRI sites of the pMT/V5-His vector (Invitrogen). In-frame Tag-RFP-T gene was then introduced at the 3’ end of γ-tubulin gene between 5’ EcoRI and 3’ NotI sites. The *Drosophila* DGT5 gene (augmin complex subunit, CG 8828) was PCR amplified from cDNA clone LD 47477 with a 5’ SpeI site and a 3’ EcoRI site. The resulting PCR product was then inserted into the 5’ SpeI and 3’ EcoRI sites of the pMT/V5 His-B vector (Invitrogen) containing in-frame mTurquoise2 gene at the 5’ end; cloned between 5’ NcoI and 3’ SpeI sites; and the Mis12 promoter at the 5’ end, cloned between a single Kpn1 site.

### Cell culture and generation of stable cell line

*Drosophila S2* cells were grown in Schneider’s medium (Life Technologies) supplemented with 10% heat inactivated fetal bovine serum (FBS) and 0.5x antibiotic/antimycotic cocktail (Sigma), and maintained at 25°C. Cell lines were generated by transfecting the DNA constructs containing the gene of interest with Effectene transfection reagent (Qiagen, Hilden, Germany), following the manufacturer’s protocol. 4 days after transfection, expression of EGFP/Tag-RFP-T/mTurquoise2-tagged proteins was checked by fluorescence microscopy. To make a stable cell line, cells were selected in the presence of Blasticidin S HCl (Thermo Fisher Scientific, Waltham, MA) and/or Hygromycin (Sigma-Aldrich) until there was no observable cell death. Thereafter, cell lines were either frozen down or maintained in the S2 media @ 25 °C without Blasticidin and/or Hygromycin B.

### RNA interference (RNAi) experiments

Around 500 base pairs of DNA template containing T7 promoter sequence at the 5’ end for D-TPX2 (also known as Ssp1/mei-38 in *Drosophila*, CG14781) was generated by PCR. Double-stranded RNA (dsRNA) was synthesized from D-TPX2 DNA template (cDNA clone RE11134) containing T7 promoter at the 5’ end at 37 °C using the T7 RiboMax Express Large-Scale RNA Production System (Promega Corp., Madison, WI), following the manufacturer’s protocol. For RNAi experiments, cells at about 25% confluency were incubated in a 35 x 10 mm tissue culture dish for an hour. Thereafter, media was carefully aspirated off the dish and 1 ml of serum-free Schneider’s media containing 20 *μ*g of dsRNA was added to the dish. After 1 hour, 1 ml of fresh media containing FBS was added to the dish and incubated for 4 days at 25 °C. Depletion of endogenous D-TPX2 was confirmed by western blots.

### Western blotting

Equal amounts of proteins were loaded into an 8% SDS-PAGE gel. After running the gel, proteins were transferred to a nitrocellulose membrane using the Trans-Blot Turbo transfer system (Bio-Rad Laboratories, Inc., Hercules, CA) for 10 minutes. Subsequently, the membrane was incubated in 5% milk (W/V made in Tris-buffered saline with 0.1% Tween (TBS-T)) for one hour. Following block, the membrane was incubated with either α-D-TPX2 antibody at 1:2000 dilution or α-tubulin (DM1-A, Sigma) antibody at 1:10,000 dilution for 1 hour, followed by 3X, 5 minute washes in TBS-T and secondary antibody incubation at 1:5000 dilution for 1 hour. Following secondary antibody incubation, membrane was washed 3X, 5 minutes in TBS-T and developed with ECL reagent (Millipore). The blot was imaged with a G:BOX system controlled by GeneSnap software (Syngene, Cambridge, U.K.). Images were further quantified to estimate the knock down efficiency with the help of Fiji/Image J software (*Schindelin et al., 2012*). To obtain the intensity values, identical regions were drawn over each band and their integrated intensity was recorded. Intensity values were normalized to their respective loading controls (tubulin) to estimate the knockdown efficiency.

### Live-cell TIRF microscopy

Cells at about 50% confluency, expressing the gene of interest were seeded on a 35mm glass bottom dish (Cellvis, CA, USA) coated with Concanavalin A for 30-60 minutes. Before imaging, the total volume was brought up to 2 ml with fresh Schneider’s media containing FBS. Multi-color, live-cell TIRF movies of all the constructs (EGFP-α-tubulin and γ-tubulin-Tag-RFP-T; EGFP-α-tubulin, γ-tubulin-Tag-RFP-T, and mTurquoise2-Dgt5) were acquired on a Nikon Ti-E microscope equipped with a 100X 1.4 NA DIC Apo Oil immersion objective, a Hamamatsu ORCA-Flash 4.0 LT digital CMOS camera (C11440), 4 laser lines (447 nm, 488 nm, 561 nm, 641 nm), and MetaMorph software. Metamorph software (Molecular Devices) was used to control the imaging systems.

## ACKNOWLEDGMENTS

We acknowledge Patricia Wadsworth (UMass, Amherst) for sharing many insightful conversations. We would like to thank Gohta Goshima (Nagoya University, Nagoya, Japan) for sharing the α-D-TPX2 (Ssp1/Mei-38) serum. We acknowledge Ted Salmon for inspiring this work and for consistently demonstrating the power of observation and quantification of cellular phenomena using high resolution light microscopy. This work was supported by a National Institutes of Health grant (GM107026) to T.J.M.

---

**Supplementary Figure 1.**
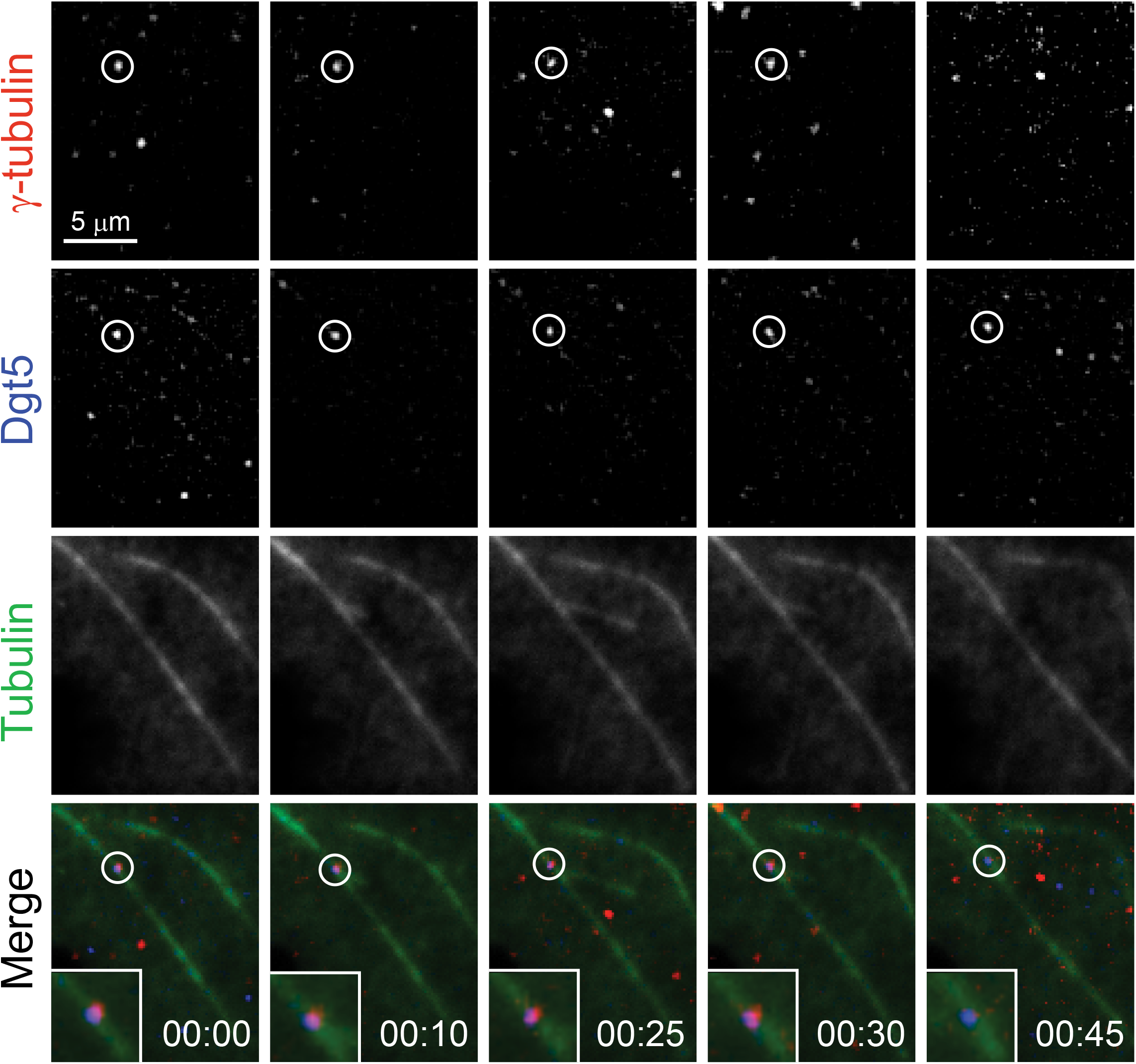
A MT branching event during anaphase in a cell co-expressing GFP-α-tubulin (green), mTurquoise2-Dgt5 (blue), and γ-tubulin-TagRFP-T (red). Co-localized Dgt5 and γ-tubulin (timepoint 00:00) nucleates a MT branching event at 00:10. Subsequent frames show polymerization (00:10 - 00:25) and depolymerization (00:30 - 00:45) of the daughter MT. Time: mins:secs. Scale bar, 5 μm.

## SUPPLEMENTAL MOVIES

**Movie 1: Direct observation of γ-tubulin-mediated MT branching.** Time-lapse TIRF microscopy showing a γ-tubulin-mediated MT branching event in an anaphase *Drosophila* S2 cell co-expressing EGFP-α-tubulin (green) and γ-tubulin-Tag-RFP-T (red). The mother MT and daughter MT both contact the plasma membrane. In the montage, γ-tubulin-Tag-RFP-T is in the upper left panel, EGFP-α-tubulin is in the upper right panel, and the merge is in the lower left panel. Time: mins:secs. Scale bar, 1 μm

**Movie 2: Direct visualization of augmin-mediated MT branching.** Time-lapse TIRF microscopy showing a MT branching event in an anaphase *Drosophila S2* cell co-expressing EGFP-α-tubulin (green), γ-tubulin-Tag-RFP-T (red), and mTurquoise2-Dgt5 (blue). In the montage, γ-tubulin-Tag-RFP-T is in the upper left panel, mTurquoise2-Dgt5 is in the upper right panel, EGFP-α-tubulin is in the lower left panel, and the merge is in the lower right panel. Time: mins:secs. Scale bar, 1 μm

**Movie 3: Direct visualization of augmin-mediated MT branching.** Time-lapse TIRF microscopy showing two microtubule branching events in a cell co-expressing EGFP-α-tubulin (green), γ-tubulin-Tag-RFP-T (red), and mTurquoise2-Dgt5 (blue) in an anaphase *Drosophila S2* cell. Two Dgt5 puncta bind to a mother MT (one indicated by an arrow 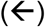 and another by a left caret (<) in the Dgt5 channel). One Dgt5 puncta (<) recruits a γ-tubulin (left caret (<) in RFP channel, indicated by 1) at 00:20; however, this γ-tubulin dissociates from Dgt5 at 00:50 without nucleating a daughter. The second Dgt5 puncta 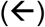 recruits γ-tubulin at 00:50 (indicated by an arrow 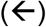 and designated 2) and nucleates a daughter MT at 01:20. The first Dgt5 puncta (<) remains localized to the mother and recruits another γ-tubulin (left caret (<) in RFP channel, indicated by 3) at 01:30, which nucleates a daughter MT at 01:40 shortly before the mother MT depolymerizes (01:50 - 02:10). In the montage, γ-tubulin-Tag-RFP-T is in the upper left panel, mTurquoise2-Dgt5 is in the upper right panel, EGFP-α-tubulin is in the lower left panel, and the merge is in the lower right panel. Time: mins:secs. Scale bar, 1 μm

**Movie 4: Branched MT arrays amplify RhoA activation near the cortex.** Time-lapse TIRF microscopy of an anaphase *Drosophila* S2 cell co-expressing Tag-RFP-T-α-tubulin and Rhotekin-GFP (a marker for active RhoA) activation near the branched MT arrays in a in *Drosophila S2* cells. Enriched Rhotekin signal is observed near the branched MT arrays; while the nearby region devoid of MT plus-ends does not exhibit Rhotekin enrichment. In the montage, γ-tubulin-Tag-RFP-T is in the upper left panel, Rhotekin-EGFP is in the upper right panel, and the merge is in the lower left panel. Time: mins:secs. Scale bar, 5 μm.

